# Skin Delivery of Modified Vaccinia Ankara Viral Vectors Generates Superior T Cell Immunity Against a Respiratory Viral Challenge

**DOI:** 10.1101/2020.05.06.079046

**Authors:** Youdong Pan, Luzheng Liu, Tian Tian, Jingxia Zhao, Chang Ook Park, Serena Y. Lofftus, Claire A. Stingley, Shenglin Mei, Xing Liu, Thomas S. Kupper

## Abstract

Modified Vaccinia Ankara (MVA) was recently approved as a Smallpox vaccine. Transmission of Variola is by respiratory droplets, and MVA delivered by skin scarification (s.s.) protected mice far more effectively against lethal respiratory challenge with VACV than any other route of delivery, and at much lower doses. Comparisons of s.s. with intradermal, subcutaneous or intramuscular routes showed that MVA_OVA_ s.s.-generated T cells were both more abundant and transcriptionally distinct. MVA_OVA_ s.s. produced greater numbers of lung Ova-specific CD8^+^ T_RM_ and was superior in protecting mice against lethal VACV_OVA_ respiratory challenge. Nearly as many lung T_RM_ were generated with MVA_OVA_ s.s. compared to direct pulmonary immunization with MVA_OVA_, and both routes vaccination protected mice against lethal pulmonary challenge with VACV_OVA_. Strikingly, MVA_OVA_ s.s.-generated effector T cells exhibited overlapping gene transcriptional profiles to those generated via direct pulmonary immunization. Overall, our data suggest that heterologous MVA vectors delivered via s.s. are uniquely well-suited as vaccine vectors for respiratory pathogens like COVID-19. In addition, MVA delivered via s.s. could represent a more effective dose-sparing smallpox vaccine.

---

Vaccines against viral and bacterial pathogens have become a fundamental part of pediatric and adult patient care^1–4^. Once ubiquitous diseases like smallpox, polio, measles, tetanus, and diphtheria have either been eliminated or substantially reduced in incidence by vaccination in most of the industrialized world. Vaccination against seasonal influenza has been more challenging, and vaccination against HIV has proven elusive^5–7^. Vaccines against emerging diseases like Ebola, SARS, and MERS and most recently COVID-19 are the subject of intense interest and widespread activity^8–10^. Most vaccines are administered by intramuscular or subcutaneous injection. While readily accessible, skeletal muscle tissue is poorly adapted to initiating immune responses, as is subcutaneous adipose tissue^11^. In contrast, upper layers of the skin are the site of continuous and multiple immune responses over a lifetime^12^. Smallpox vaccination through skin with Vaccinia virus (VACV) has been uniquely successful^2,11^.

The eradication of smallpox by worldwide epicutaneous immunization with VACV was the greatest public health achievement of the 20^th^ century^2^. Since that time, VACV has been employed as a vaccine vector in many settings^13^. However, its use has been limited by unacceptable morbidity, particularly in recipients who are immunocompromised^14^. More recently, Modified Vaccinia Ankara (MVA), a replication-deficient variant of VACV, has come into wider use^15^. Although it lacks ∼10% of the parent genome^16^, it retains the immunogenicity of the parent virus and has just been approved by the FDA as a modern alternative for smallpox preventative vaccination^17^. Like VACV, it is also being widely used as heterologous vaccine vector^18^. However, MVA and derivative vectors are almost invariably delivered intramuscularly or subcutaneously^19^.

Several important features of smallpox vaccination deserve to be re-emphasized. In the 20^th^ century, delivery of VACV i.m. was ineffective at conferring protection against smallpox. In contrast, development of a cutaneous “pox” lesions, achieved only after epicutaneous immunization, was considered emblematic of successful protective vaccination, suggesting that this mode of delivery was critically important^14^. In addition, smallpox vaccination was effective in patients with agammagloblulinema, while VACV immunization had disastrous complications in patients with T cell deficiency^20^. This suggested that T cells were critically important for protective immunity^21,22^. Finally, Variola virus is transmitted via respiratory droplets, suggesting an oropharyngeal-pulmonary mode of transmission^23^. It is notable that murine models of epicutaneous skin immunization with VACV generate memory T cell populations in both skin and lung, and these lung memory T cells protect against lethal pulmonary challenge with this virus^11^. Intramuscular immunization with VACV in these models did not yield comparable protection. This suggests that protection against smallpox is at least in part mediated by T cells^22,24^, and that skin immunization is an effective means of generating protective memory T cell populations in the lung^11^.

In the present study, we asked whether immunization with MVA was more effective and more effective if delivered epicutaneously (s.s.) as compared to intramuscularly (i.m.). We also asked whether skin immunization with an MVA vector generated populations of antigen specific CD8^+^ T cells in lung as well as skin. In addition, epicutaneous immunization was compared to intradermal, subcutaneous, and intramuscular immunization in generated protective immunity against a lethal pulmonary challenge. Finally, we asked if T cell imprinting by skin draining and lung draining nodes was similar.

## Results

Doses from 10^4^ pfu to 10^7^ pfu of MVA were used for epicutaneous immunization (s.s.), and after 7 days, lymph node and spleen T cells were harvested and stimulated *in vitro* with VACV infected splenocytes, after which IFN-γ production was measured. All MVA doses led to significant T cell IFN-γ production, with 10^6^ and 10^7^ pfu being equally potent **(Fig. 1a,b)**. Other groups of mice were immunized with these doses, and after 30 days these mice were challenged on the skin with VACV. After 6 days, biopsies of the immunized sites were taken and VACV DNA was measured by PCR. All immunization doses led to diminished VACV DNA at the infected site (compared to unimmunized controls), but 10^6^ and 10^7^ pfu immunization showed superior protection **(Fig. 1c)**. Other groups of mice were immunized in an identical manner and were subjected to lethal intranasal infection with VACV at day 30. All unimmunized mice rapidly lost weight and succumbed to the infection. In contrast, 40% of 10^4^, 70% of 10^5^, and 100% of 10^6^ and 10^7^ pfu immunized mice survived the infection **(Fig. 1d,e)**. Thus, 10^6^ pfu is the lowest MVA dose that provides both strong T cell cytokine production as well as optimal protective immunity against skin and pulmonary infection.

**Figure 1.**
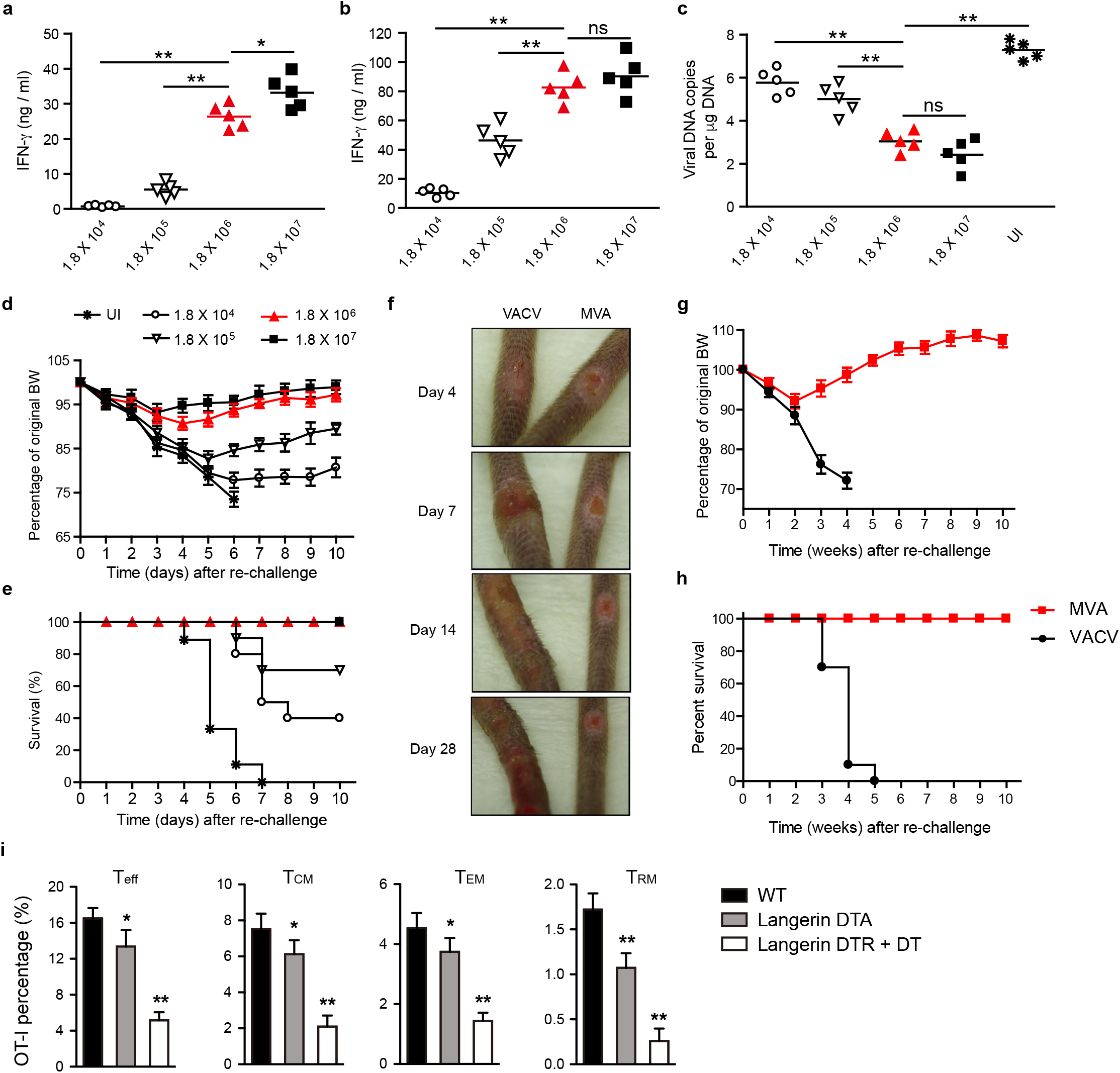
MVA immunization via skin scarification (s.s.) elicits dose-dependent anti-vaccinia immune response. **a-b.** IFN-γ secretion by vaccinia-specific T cells isolated from draining lymph nodes (a) or spleens (b) at 7 days post MVA infection at indicated dose. **c.** Quantitative real time PCR (qRT-PCR) analysis of skin viral load at 6 days post re-infection. Mice were immunized with the indicated doses of MVA via s.s. 45 days later, mice were rechallenged with 1.8 × 10^6^ pfu vaccinia virus (VACV). Then 6 days later, skin tissues were harvested and processed to qRT-PCR. **d-e.** Body weight (BW) (d) and survival measurements (e) of WR-VACV re-challenged mice that were immunized previously with MVA at indicated dose 45 days earlier. **f.** Photographs of pox lesion in Rag1^-/-^ mice taken on day 4, 7, 14 and 28 postimmunization with the same amount (1.8 × 10^6^ pfu) of MVA or VACV. **g-h.** Immunized Rag1^-/-^ mice were monitored for BW change (g) and survival (h) for up to 12 weeks after immunization with the same amount (1.8 × 10^6^ pfu) of MVA or VACV. **i.** Quantification of effector T cell (T_eff_, day 5), central memory (T_CM_, day 45), effector memory (T_EM_, day 45) or tissue resident memory (T_RM_, day 45) T cells post MVA infection. Naïve OT-I Thy1.1^+^ cells were transferred into Thy1.2^+^ recipient mice one day before mice were infected with 1.8 × 10^6^ pfu MVA-Ova. Then at different time points post infection, OT-I cells were isolated from lymph nodes (T_eff_, T_CM_, T_EM_) or skin (T_RM_) and analyzed by flow cytometry. a-c. Data is representative of three independent experiments. Symbols represent individual mice (n = 5 mice/group). c-d. Unimmunized (UI) mice were included as controls. Graphs show mean ± s. d., ns = not significant, *p < 0.05, **p < 0.01.

To test whether delivery of MVA to scarified skin could induce poxvirus-specific immune responses, we inoculated C57BL/6 mice with MVA or Vaccinia Virus (VACV) by scarification. By 7 days after inoculation, a pustular lesion resembling a “pox” reaction had formed at the inoculation site in all the immunized mice. The pox lesions induced by MVA and VACV skin scarification followed similar patterns of evolution (although with different size and kinetics), from macules to papules to vesicles and finally into pustules which ruptured and healed over time with scars **(Extended data, Fig. 1)**. MVA-induced pox reactions did heal more rapidly than those induced by replication competent VACV **(Extended data, Fig. 1)**. To determine the safety of MVA in immunocompromised hosts, we next immunized immunodeficient Rag1^-/-^ mice with VACV and MVA, respectively, and followed the mice for several weeks. While both groups of mice lost some weight over the first two weeks, MVA immunized mice rapidly regained the weight and flourished over the next several weeks **(Fig. 1g)**. In contrast, 100% of the VACV immunized mice developed progressive weight loss and expanding cutaneous lesions of VACV infection, ultimately requiring euthanasia **(Fig. 1f-h)**. Thus, MVA can be administered safely to mice wholly deficient in adaptive immunity. In another set of experiments, we immunized Wild-type (WT) mice as well as mice deficient in either Langerhans cells (Langerin DTA) or both Langerhans cells and langerin positive dermal dendritic cells (Langerin DTR + DT), respectively. Prior to infection, mice were loaded with OT-1 cells and the immunizing virus was MVA_OVA_. Spleen and lymph nodes were harvested at days 10 and 30, skin was harvested at day 30, and OT-1 T cells were counted. At day 10, T_eff_ cells were somewhat diminished in Langerin DTA mice and more markedly diminished in Langerin DTR + DT mice **(Fig. 1i)**. At day 30, skin T_RM_ were significantly diminished in Langerin DTA mice and even more diminished in Langerin DTR+DT mice **(Fig. 1i)**. This pattern was also true for T cells bearing markers of T_CM_ and T_EM_. These data suggest that both LC and langerin positive dermal DC play an additive role in optimal antigen presentation of MVA-encoded antigens to T cells.

We next compared the anatomical route of vaccine delivery on the T cell response to MVA vaccination. Using CFSE OT-1 loaded mice, MVA_OVA_ was delivered by epicutaneous infection (s.s.), or injected intradermally (i.d.), subcutaneously (s.c.), or intramuscularly (i.m.). Draining lymph nodes were harvested at 60 hours and 5 days, and OT-1 cells were analyzed by FACS. LN from s.s. immunized mice showed roughly 90% of OT-1 proliferating, and 60% making IFN-γ, at 60 hours, with comparable numbers at 5 days **(Fig. 2a,c)**. Vaccination by i.d. was less effective, with 71% of OT-1 cells proliferating and 33% making IFN-γ at 60 hours, with modest improvement at 5 days post infection **(Fig. 2a,c)**. Both s.c. and i.m. showed poor OT-1 activation at 60 hours with some improvement at 5 days **(Fig. 2a,c)**. When lymph node or spleen OT-1 cells were stimulated with antigen, significantly more IFN-γ was produced by OT-1 cells from mice vaccinated via s.s. compared to other routes **(Extended data, Fig. 2)**. Vaccination via i.d. was intermediate with regard to IFN-γ production, while s.c. and i.m. led to nearly four-fold lower IFN-γ levels **(Extended data, Fig. 2)**. In terms of absolute numbers of OT-1 cells generated, s.s. was superior to all modes of vaccination, with i.d. being second and both i.m. and s.c. far less effective **(Fig. 2b,d)**. We next took OT-1 cells from the 5-day post-immunization time point and performed transcriptional profiling on OT-1 cells generated after s.s., i.d., s.c., or i.m., respectively. While there was some overlap, there were surprisingly many differences between T cells generated by different routes of immunization, even at the same day post immunization **(Fig. 2e, Extended data, Fig. 3)**. Principal component analysis revealed that T_eff_ generated by s.s. and i.m. were transcriptionally quite distinct. T cells generated after s.s., i.d., and s.c., were more similar but still quite distinct from one another. T cells generated by s.s. and i.d. clustered closely but were still clearly not overlapping. Moreover, s.s generated most abundant skin infiltrating cells at day 5 post immunization **(Fig. 2g)**.

**Figure 2.**
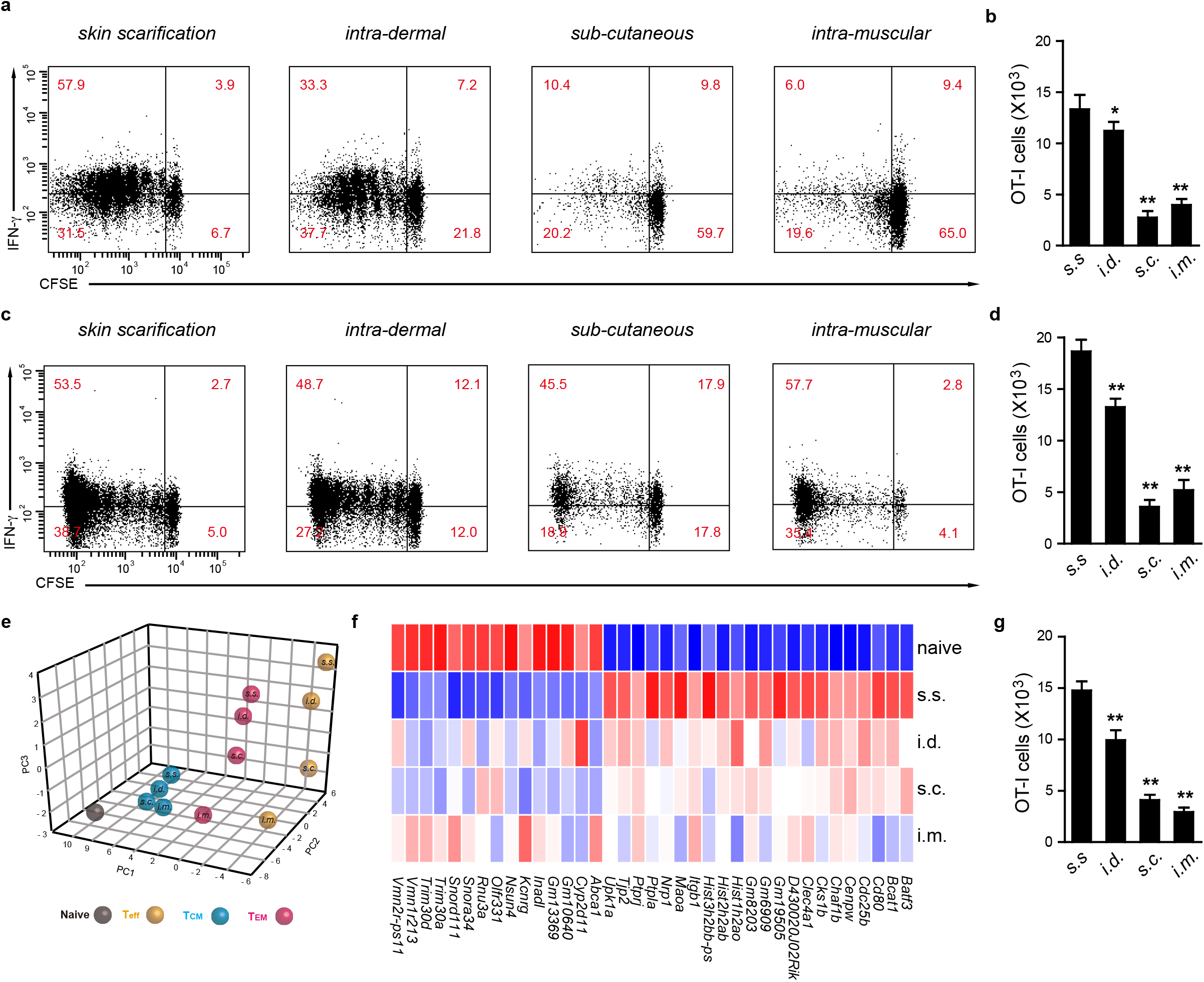
Delivery of MVA via s.s. generates T cells that are both quantitatively more abundant and qualitatively distinct from those generated from i.d., s.c., i.m.. **a-d.** Flow cytometric analysis (a, c) and quantification (b, d) of OT-I cell proliferation in draining lymph nodes of recipient mice at 60 hours (a, b) and 5 days (c, d) post MVA infection via different routes. CFSE-labeled naïve OT-I Thy1.1^+^ cells were transferred into Thy1.2^+^ recipient mice one day before mice were infected with 1.8 × 10^6^ pfu MVA-Ova via indicated infection routes. **e**. Principal component analysis (PCA) of gene-expression for T cells generated by MVA infection via different routes. Naïve T cells (T_N_) were sorted from the peripheral lymph nodes of naïve OT-I mice. Effector T cells (T_eff_) were sorted from draining lymph nodes at 5 days post infection. Central memory T cells (T_CM_) and effector memory T cells (T_EM_) were sorted from the spleen of mice at 45 days post infection. Each dot represents an individual experiment wherein mRNA was pooled from 15-20 mice from 3-4 independent biological groups (5 mice/group). **f.** Heatmap of differentially expressed genes selected from a pair-wise comparison between s.s. generated T_eff_ cells and naïve T cells. **g.** Quantification of skin infiltrating T cells at day 5 post 1.8 × 10^6^ pfu MVA-Ova infection via indicated routes. a, c, Data are representative of three independent experiments (n = 5 mice per group). Graphs show mean ± s. d. of 5 mice per group. *p < 0.05, **p < 0.01.

We next examined memory OT-1 T cells generated at 45 days by these four routes of immunization. With regard to T_CM_, s.s. generated the largest population of these cells, roughly twice as many as i.m. **(Fig. 3a)**. The difference was even more striking when T_EM_ were examined; here, s.s. generated at least 3-fold more cells than did other modes of immunization, with s.c. being least effective **(Fig. 3b)**. T_RM_ were then examined, in both skin and lung. Immunization via s.s. generated 3-fold more skin T_RM_, and more than twice as many T_EM_, with i.d. being the second most effective route **(Fig. 3-f)**. Because MVA is often delivered i.m., it is important to note that the number of T_RM_ generated by this route was more than 4-fold lower than by s.s. **(Fig. 3 c-f)**. Transcriptional profiling showed that at 45 days, OT-1 T_EM_’s still showed non-overlapping PCA clusters from s.s, i.d., s.c., and i.m. immunized mice. In contrast, T_CM_ from the same mice showed transcriptional profiles that were more tightly clustered, indicating that differences between the groups were minimal **(Fig. 2e)**. Skin T_RM_ could not be compared because insufficient 45-day T_RM_ were generated by i.m. and s.c. immunization.

**Figure 3.**
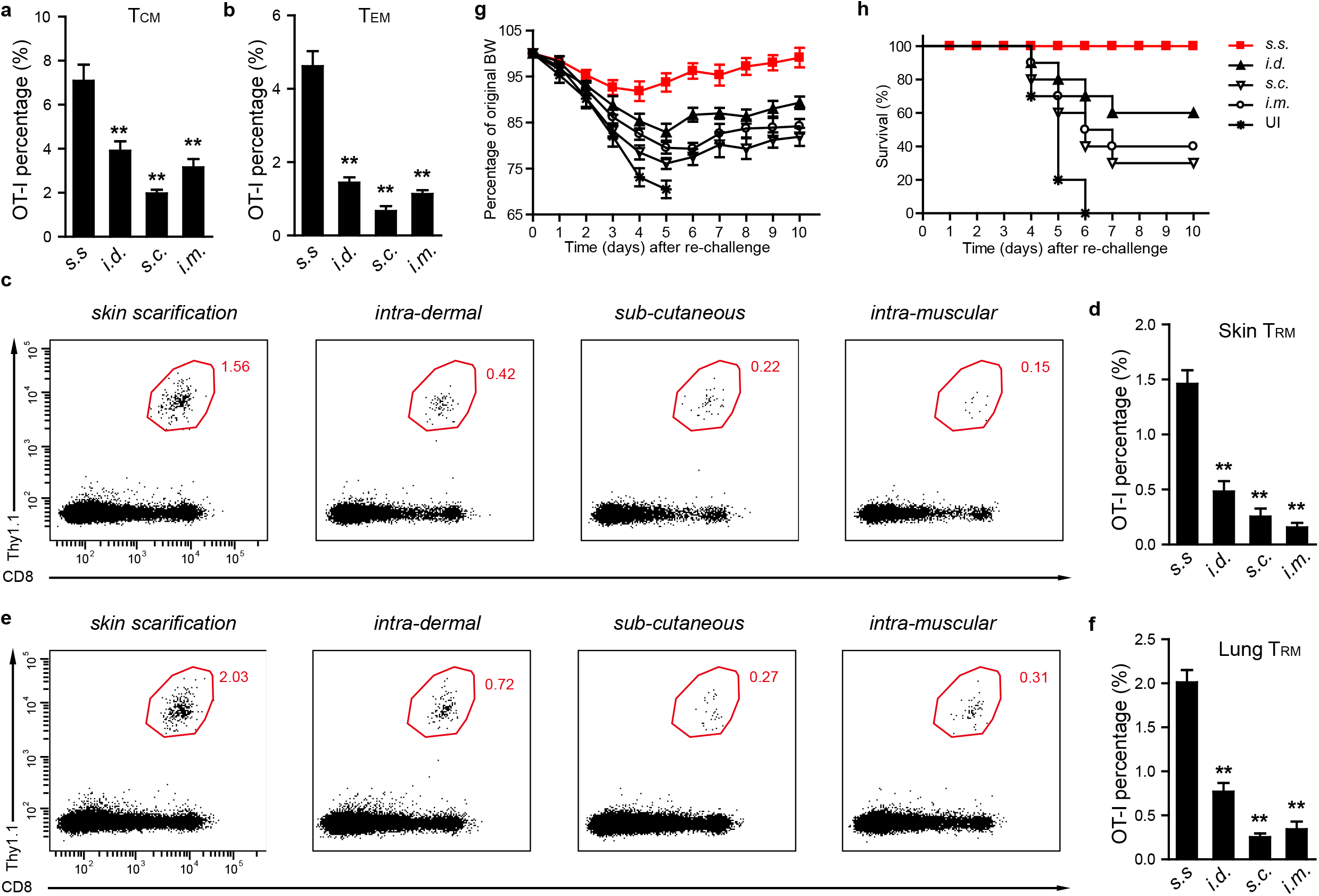
Delivery of MVA via s.s. is superior in generating memory T cells and is superior in protecting mice against lethal respiratory challenge. **a-b.** Quantification of OT-I T_CM_ and T_EM_ cells from spleen of mice at 45 days post MVA infection via indicated routes. **c-f.** Flow cytometric analysis (c, e) and quantification (d, f) of OT-I T_RM_ cells isolated from skin (c, d) or lung (e, f) tissue at 45 days post MVA infection via indicated routes. **g-h.** Body weight (BW) (g) and survival measurements (h) of WR-VACV re-challenged mice that were previously immunized with MVA via indicated routes 45 days earlier. OT-I WT cells were adoptively transferred into μMT mice before mice were infected with 1.8 × 10^6^ pfu MVA via indicated routes. 45 days later, mice were re-challenged with a lethal dose of WR-VACV by intranasal infection. c, e, Data are representative of three independent experiments (n = 5 mice per group). Graphs show the mean ± s. d. of 5 mice per group. Un-immun. = un-immunized. **p < 0.01.

In subsequent experiments, we examined groups of mice vaccinated by these different routes for their ability to respond to a lethal intranasal challenge of VACV_OVA_. Groups of ten mice assayed 45 days after initial vaccination were subjected to intranasal challenge, and mice were weighed daily after vaccination. Mice that lost >20% of body weight were sacrificed. **Figure 3g** and **h** show that naïve mice universally succumbed to the lethal infection, while mice immunized epicutaneously (s.s.) showed minor transient weight loss but complete survival. In contrast, mice vaccinated i.d., s.c., or i.m. lost substantial weight **(Fig. 3g)**, and while 60% of i.d. vaccinated mice survived, only 40% and 30% of mice vaccinated i.m. and s.c., respectively, survived **(Fig. 3h)**. These results are consistent with the superior production of different memory T cell subsets after vaccination by s.s..

We were struck by the capacity of skin immunization via s.s. to generate both skin T_RM_ and lung T_RM_. While skin and gut T cell trafficking have been studied extensively, lung T cell trafficking has been studied less comprehensively. We immunized CFSE OT-1 loaded mice with MVA_OVA_ via three routes: s.s. to assess skin homing, intraperitoneally (i.p.) to assess gut homing, and intra-tracheally (i.t.) to assess lung homing. At 60 hours, T cells were collected from the respective draining lymph nodes (inguinal for skin, mesenteric for gut, and mediastinal for lung) and were sorted based on CFSE expression into cells that had not divided (P0) or had divided once through five times (P1-P5; **Fig. 4a**). Cells were subjected to transcriptional profiling, and results were analyzed bioinformatically. By principal component analysis, P0 cells from skin, gut, and lung homing nodes clustered near each other **(Fig. 4b)**. However, as early as P1 and clearly by P2, OT-1 cells activated in different nodes diverged significantly in transcriptional profile. In particular, OT-1 cells from mesenteric nodes were quite distinct from OT-1 cells from inguinal and mediastinal nodes **(Fig. 4b)**. Interestingly, P1-P5 cells from inguinal (skin draining) node clustered closely with P1-P5 cells from mediastinal (lung draining) nodes, suggesting similar pathways involved in skin and lung homing imprinting **(Fig. 4b)**. Excluding genes upregulated in all T cell groups, lung and skin homing T cells shared upregulation of 150 genes, compared to 73 and 90 upregulated in only skin or only lung, respectively **(Fig. 4c, d)**. In contrast, only 11 upregulated genes were shared between skin and gut, and only 36 between lung and gut. Examination of chemokine receptors and integrin genes showed homology between lung and skin, while gut immunization showed unique upregulation of CCR9, α4 and β7 integrins **(Fig. 4e)**. These data suggest a very similar pattern of gene expression of T cells activated in skin versus lung draining LN, and a pattern in gut draining LN that is very different from lung and skin draining LN.

**Figure 4.**
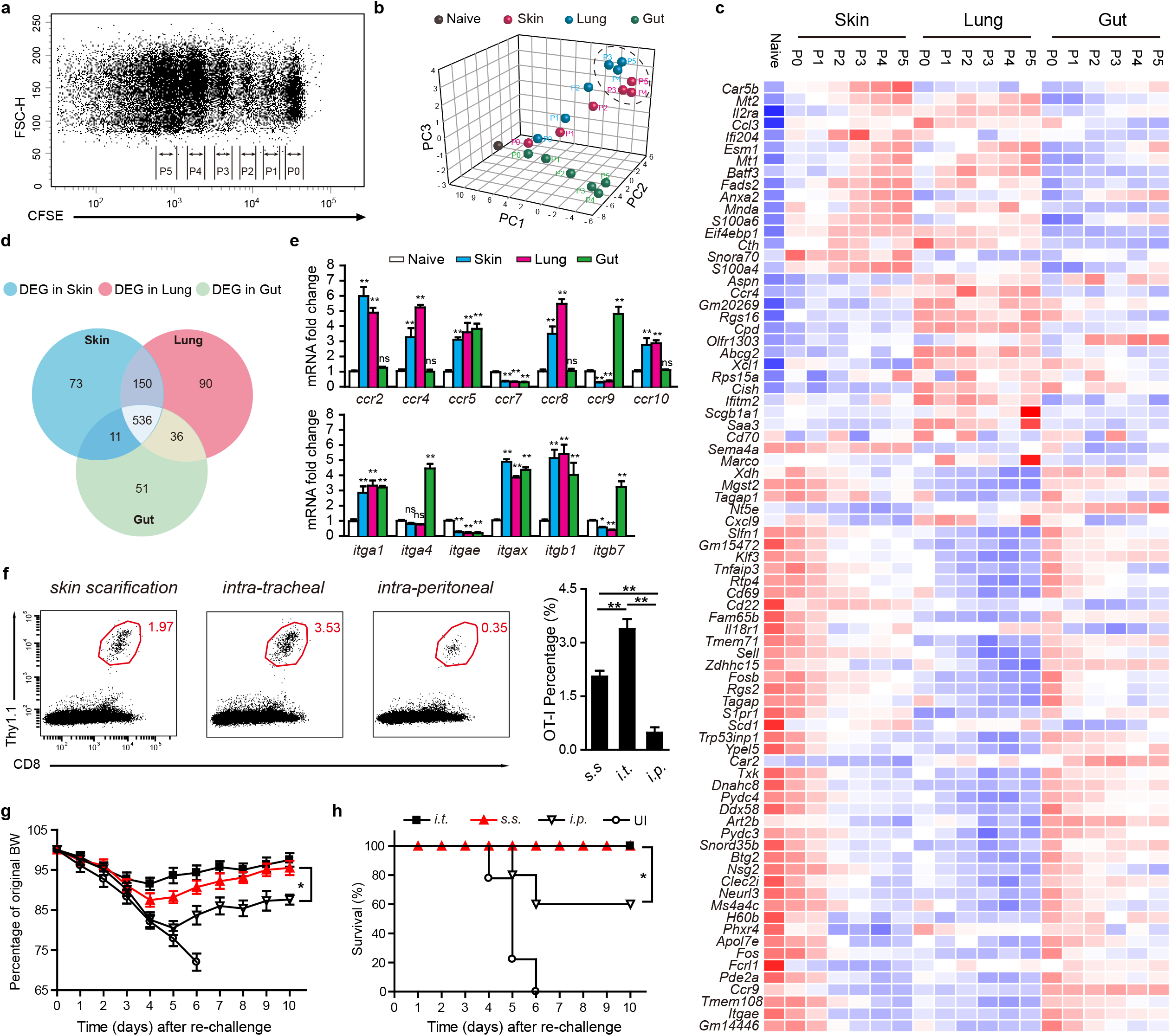
MVA s.s. generates more than half number of lung T_RM_ compared to intratracheal (i.t.) and is sufficient to protect mice against lethal respiratory challenge. **a.** Flow cytometric analysis of OT-I cell proliferation in draining lymph nodes at 60 hours post MVA infection via s.s.. CFSE-labeled naïve OT-I Thy1.1^+^ cells were transferred into Thy1.2^+^ recipient mice one day before mice were infected with 1.8 × 10^6^ pfu MVA-Ova. **b**. PCA of geneexpression data for nineteen CD8^+^ T cell populations based on CFSE signal and different infection routes. Each dot represents an individual experiment wherein mRNA was pooled from 15-20 mice from 3-4 independent biological groups (5 mice/group). **c.** Heatmap of differentially expressed genes selected from a pair-wise comparison between s.s. and intra-peritoneal (i.p.) activated T cells. **d**. Venn diagram analysis of genes differentially expressed in pairwise comparisons between s.s., i.t. and i.p. activated T cells relative to T_N_ (fold change cutoff, ≥2). **e**, Quantitative real-time PCR (qRT-PCR) analysis of cell homing molecule gene expression in s.s., i.t. and i.p. activated T cells. **f**. Flow cytometric analysis (left) and quantification (right) of lung T_RM_ cells at day 45 post MVA infection via indicated routes. **g-h.** Body weight (BW) (g) and survival measurements (h) of WR-VACV re-challenged mice that were immunized previously with MVA via indicated routes 45 days earlier. OT-I WT cells were adoptively transferred into μMT mice before mice were infected with 1.8 × 10^6^ pfu MVA via indicated routes. 45 days later, mice were re-challenged with a lethal dose of WR-VACV by intranasal infection. Graphs show the mean± s. d. of 5 mice per group. Un-immun. = un-immunized. ns = not significant, *p < 0.05, **p < 0.01.

We next directly compared the capacity of skin (s.s.), lung (i.t.), and gut (i.p.) immunization with MVA_OVA_ to generate lung T_RM_. Mice were immunized by the above routes and after 45 days, lung T_RM_ were analyzed. As expected, lung immunization resulted in the highest number of lung T_RM_, but skin immunization by s.s. generated more than half as many T_RM_ in lung **(Fig. 4f)**. In contrast, i.p. immunization resulted in less than 10% of the lung T_RM_ compared to lung immunization **(Fig. 4f)**. Like skin T_RM_, lung T_RM_ were CD69^+^, CD103^+^, CD62L^-^, KLRG1^-^, and expressed E and P selectin ligands **(Extended data, Fig. 4)**. A companion cohort of mice were subjected to lethal intranasal challenge with VACV_OVA_. Mice immunized i.t. or s.s. showed mild weight loss but 100% recovery and survival **(Fig. 4g,h)**. Mice immunized i.p. showed more severe weight loss, and only 60% survived the infectious challenge **(Fig. 4g,h)**. In another series of experiments, i.t. immunization was compared to s.s. immunization with regard to generation of skin T_RM_. While s.s. was most efficient at generating skin T_RM_, lung immunization via i.t. generated 50% of the skin T_RM_ compared to s.s. immunization **(Extended data, Fig. 5)**. These data confirm that lung immunization can generate abundant skin T_RM_, and skin immunization can generate abundant lung T_RM_.

## Discussion

Smallpox vaccination via epidermal disruption using Vaccinia virus (VACV) provided broad and effective protective immunity against Smallpox caused by Variola major, and led to the eradication of this devastating infectious disease^14^. MVA is derived from VACV but has lost 10% of the parent genome, including several immune inhibitory genes that block CC chemokines, IFNα/β, IFNγ, TNFα, and STING^25^, and does not replicate in mammalian cells^18^. In addition to its use as a smallpox vaccine, MVA has been used extensively as a heterologous vaccine vector^15^, although we were unable to find any description of it being delivered through skin scarification. Rather, intramuscular or subcutaneous injection appear to be the preferred routes. There are no clear reasons that MVA has not been delivered via s.s., other than the assumption that replication was required for this route of administration. Here, we show that MVA delivered by s.s. can provoke a potent immune response at doses much lower than those used for i.m. and s.c. injection. In a direct comparison of delivery via i.m., s.c., and i.d. routes, s.s. administration of lower doses of MVA provide superior protective immunity against a lethal VACV challenge. These data suggest that like VACV, MVA delivered by s.s. provides a potent and durable immune response. We found that both Langerhans cells and CD207+ dermal dendritic cells were both required for optimal immunization via this route. In contrast to VACV, mice deficient in adaptive immunity could be safely immunized via s.s. with MVA, supporting the safety of this vector in immunocompromised hosts. One other advantage of the s.s. mode of delivery is dose sparing—doses of MVA too low to elicit immune response i.m. are quite immunogenic when delivered by s.s.

When used as a heterologous vaccine vector encoding for a T cell antigen, MVA_OVA_ delivered s.s. provided robust early activation of OVA-specific T cells (OT-1) in skin draining lymph nodes. Interestingly, early CD8^+^ effector T cells in skin draining lymph nodes at day 5 showed different patterns of gene expression after immunization s.s., i.d., s.c., and i.m., respectively. T cells generated by i.m. immunization were most distinct transcriptionally from those generated by s.s. immunization. When T cells were harvested from spleens at day 45 after immunization, cells with T_EM_ markers retained distinct transcriptional profiles, with i.m. generated T_EM_ cells being most distinct from s.s. generated T_EM_ cells. Day 45 memory T cells expressing CD62L (T_CM_) showed smaller transcriptional differences across immunization routes, but s.s. generated T_CM_ cells were still readily distinguished from those generated by i.m. immunization. These surprising data suggest that there are qualitative differences in T_eff_ and TM cells generated by immunization route that are evident by day 5 and persist at day 45.

There were also quantitative differences in TM generation depending on route of administration. Immunization via s.s. generated greater numbers of both T_EM_ and T_CM_ at 45 days after immunization. When skin T_RM_ were measured, s.s. generated more T cells than other routes, with i.m. being least efficient. Because lethal intranasal challenge with VACV results in death from pulmonary inflammation, we also measured lung T_RM_. Strikingly, s.s. generated higher numbers of lung T_RM_ than other routes, consistent with previous reports^11,26^, with i.m. generating fewest lung T_RM_. T_RM_ from skin and lung both expressed CD69 and CD103, with expression of E- and P-selectin ligands detectable as well. When animals were challenged by lethal intranasal infection with VACV_OVA_, only mice immunized by s.s. showed minimal weight loss and 100% survival. Mice immunized by all other routes showed greater morbidity and some mortality, with i.m. immunization being least effective. Whether the ability of s.s. immunized mice to uniformly survive the intranasal challenge of VACV_OVA_ was due to higher numbers of lung T_RM_, circulating T_EM_ and T_CM_, or qualitatively different T_eff_ and memory T cells cannot be determined from these data. However, this suggests that the original method of smallpox vaccination— s.s. administration—appears to be uniquely effective at generating robust protective immunity against airway challenge.

Because s.s. immunization was so efficient at generating lung T cells and protective immunity against a pulmonary infectious challenge, we compared skin infection with direct lung infection, and assess T_eff_ in skin and lung draining LN, respectively, using i.p. injection and mesenteric nodes as a control. Thus, three routes of immunization were compared— s.s., intratracheal (i.t.), and intraperiotoneal (i.p.), and T_eff_ from draining lymph nodes—inguinal, mediastinal, and mesenteric, respectively---were compared by transcriptional profiling. While proliferating T_eff_ from skin graining and gut draining nodes rapidly diverged, proliferating T_eff_ from skin draining and lung draining nodes showed significant overlap over time. Both α1β1 intergrin, CCR4, and CCR8 were preferentially elevated in T cells from skin and lung draining nodes, and α4β7 and CCR9 were preferentially upregulated in mesenteric lymph nodes, consistent with previously reported data^27–29^. When lung T_RM_ were examined after 45 days, both skin and lung infection generated abundant lung T_RM_, while i.p. immunization was less efficient at generating these cells. Protection against lethal intranasal challenge was complete in skin and lung immunized mice, but incomplete after i.p. immunization. These data suggest that there is substantial overlap in T cells imprinted by skin and lung draining lymph nodes and suggests that skin immunization is well-suited at generating T cells with lung tropic properties.

Two important conclusions can be drawn from this study that are relevant to human disease. First, immunization with MVA generates powerful immunity, but like VACV the most potent local and systemic immunity generated occurs after superficial skin immunization (s.s.) that involves epidermal disruption. The dose of MVA used in s.s. delivery can be much lower than required in muscle/i.m. delivery. This suggests that doses of MVA being stockpiled in anticipation of a dystopian future smallpox attack may protect orders of magnitude for more people if delivered s.s. instead of MVA. The second conclusion is that MVA delivered by s.s. is a very effective way of generating protective T_RM_ in lung, in addition to a more robust circulating T cell response. MVA vaccines are being developed for respiratory pathogens, including influenza A and respiratory syncytial virus^30,31^, but these are being tested only by i.m. or s.c. injection. Our data strongly suggests that delivering these vaccines via s.s. may generate even more effective protective immunity to pathogens that infect lung. Whether MVA encoding for Coronavirus genes and delivered s.s. could provide protective immunity against COVID-19 is an intriguing question that we are pursuing presently.

## Methods

Methods and associated references are available in the online version of the paper.

## Acknowledgements

We thank Dr. Mike Seaman (Beth Israel Deaconess Medical Center, Boston, MA) for providing the initial stock of MVA, Dr. Bernard Moss (NIH) for providing the initial stocks of VACV. This work was supported by grants R01 AI127654 to Dr. T. S. Kupper from National Institutes of Health/NIAID and R01 AR065807 from the National Institutes of Health/NIAMS.

## Supplementary Information

Supplementary Information is linked to the online version of the paper.

## Extended Data Figure Legends

**Extended Data Figure 1.**
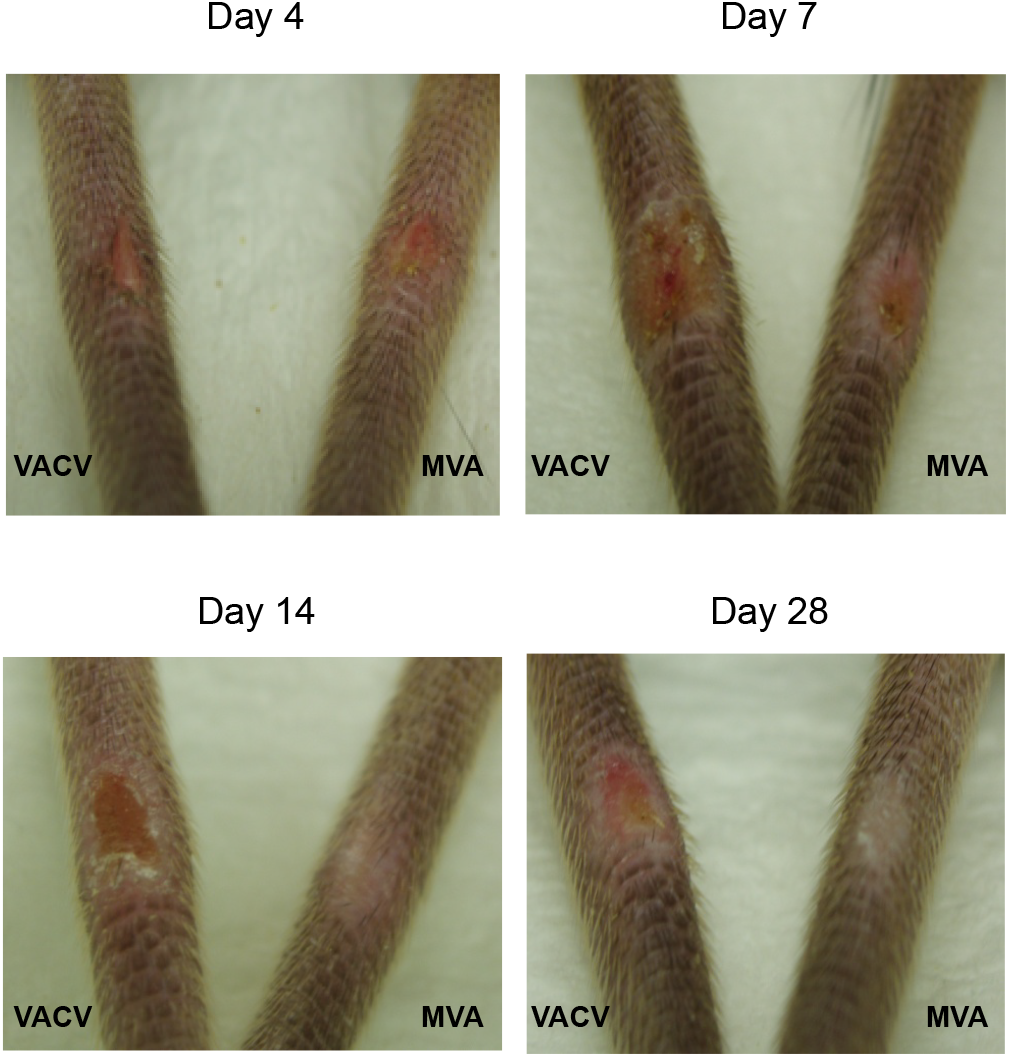
MVA skin scarification induced smaller pox lesions that healed significantly faster compared to VACV skin scarification in immunocompetent mice. C57BL/6 mice were immunized with 1.8 × 10^6^ pfu MVA or VACV by skin scarification. Photographs of pox lesion were taken on day 4, 7, 14 and 28 post-immunization.

**Extended Data Figure 2.**
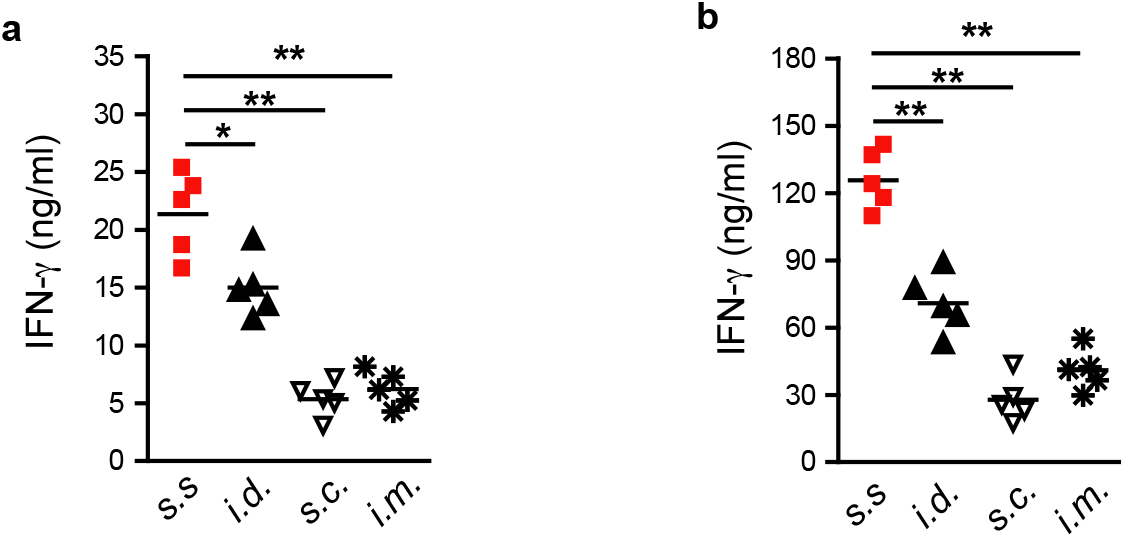
Delivery of MVA via s.s. generates stronger cellular responses compared to i.d., s.c., and i.m. infection routes. C57BL/6 mice were immunized with 1.8 × 10^6^ pfu MVA via indicated routes. Activated T cells in draining lymph nodes (a) and spleen (b) were isolated at 7 days post infection, and T cell response against VACV was measured based on IFN-γ secretion. Symbols represent individual mice (n = 5 mice/group). *p < 0.05, **p < 0.01.

**Extended Data Figure 3.**
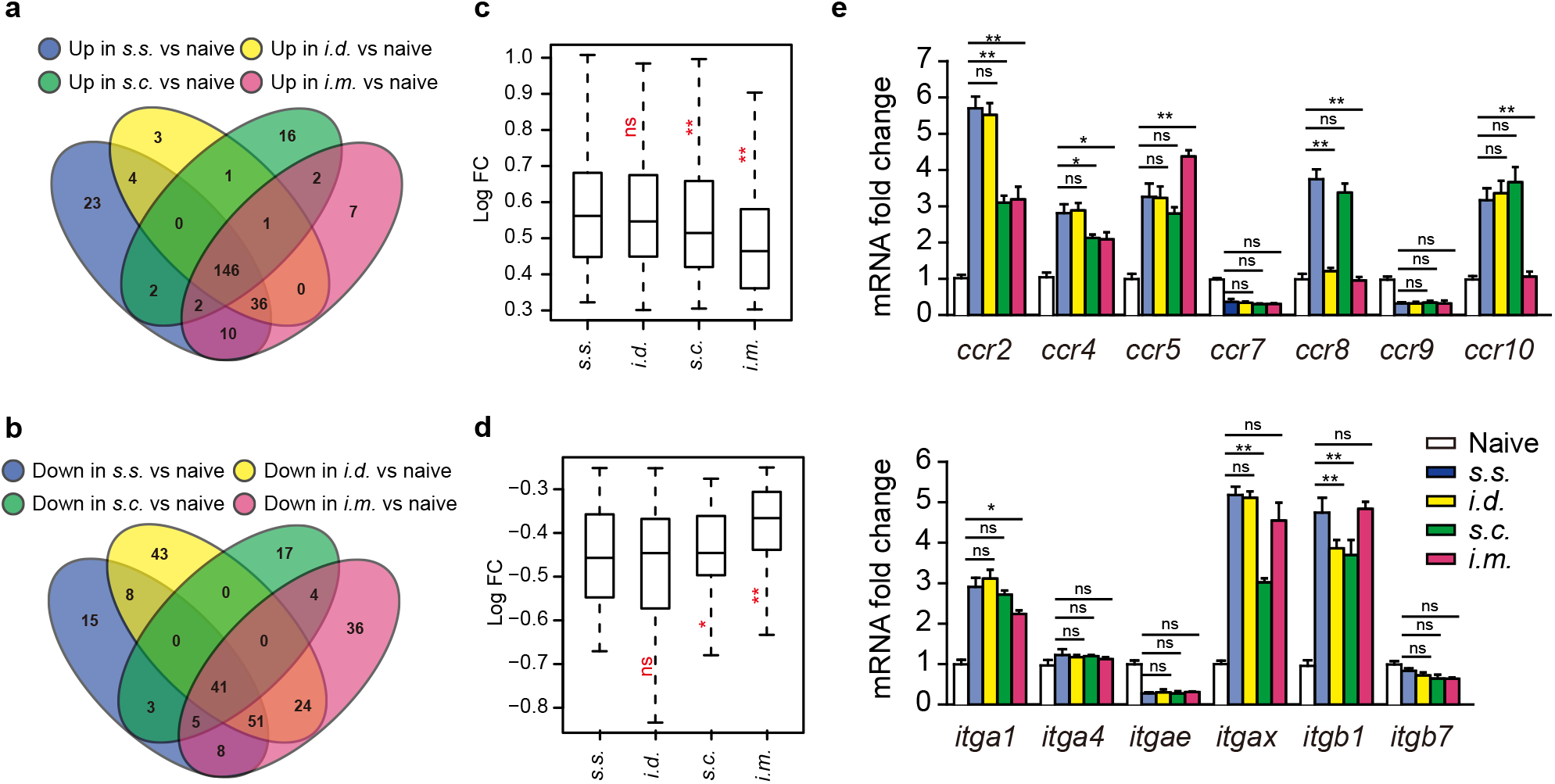
Delivery of MVA via s.s. generates T cells that are qualitatively distinct from those generated from i.d., s.c., i.m.. **a-b.** Venn diagram analysis of genes up-regulated (a) or down-regulated (b) in pairwise comparisons between T cells activated via MVA s.s., i.d., s.c., i.m. (day 5) relative to that of T_N_. **c-d.** Fold change analysis of genes shared among s.s., i.d., s.c. and i.m. activated T cells (day 5) relative to that of T_N_. c, 146 shared up-regulated genes, d, 41 shared down-regulated genes. **e.** Quantitative real-time PCR (qRT-PCR) analysis of cell homing molecule gene expression in s.s., i.d., s.c. and i.m. activated T cells (day 5) relative to that of T_N_. ns = not significant, *p < 0.05, **p < 0.01.

**Extended Data Figure 4.**
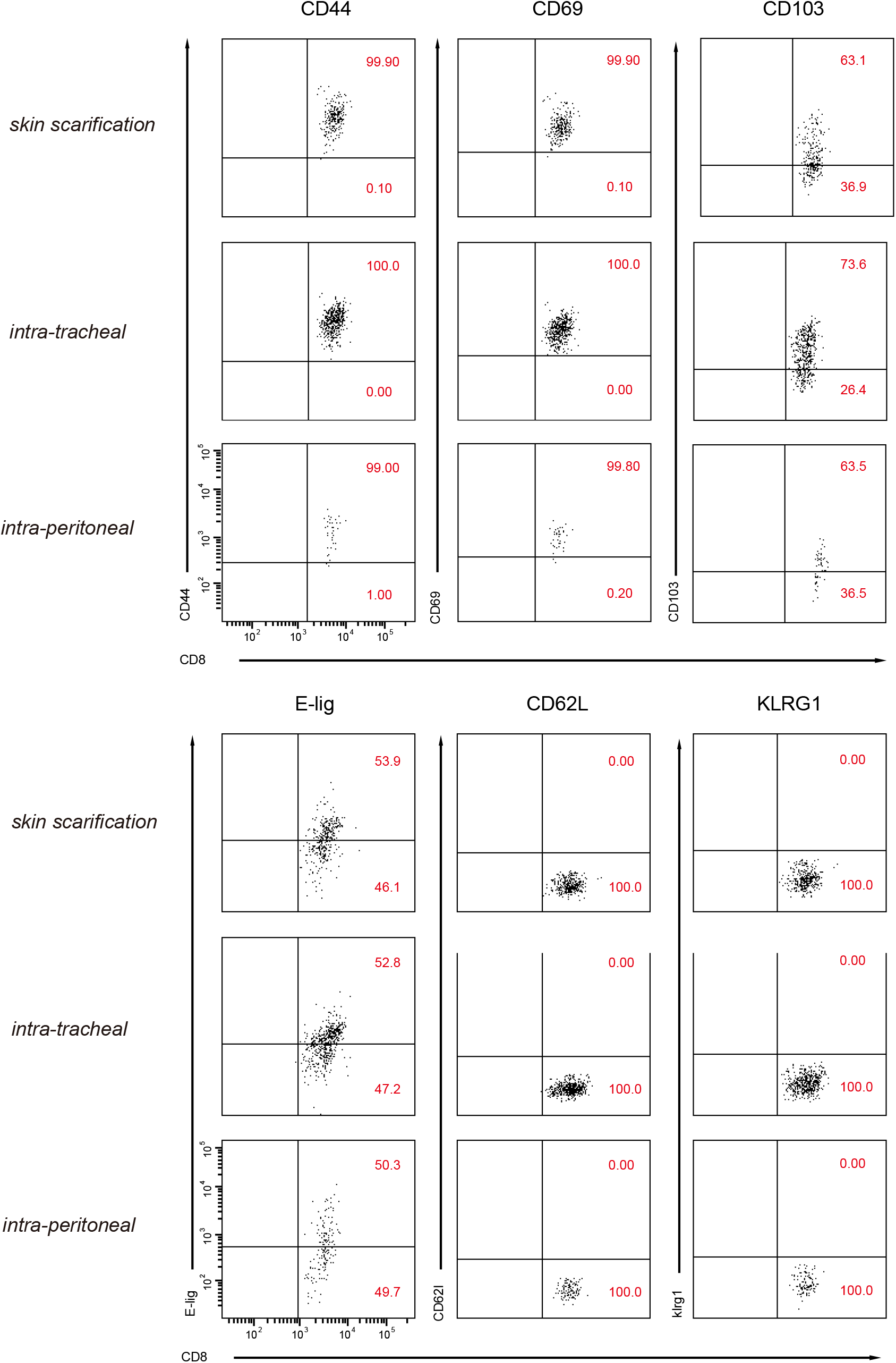
Phenotyping of tissue-resident memory T cell surface marker on lung CD8^+^ T_RM_ cells generated by MVA infection via skin scarification, intra-tracheal administration or intra-peritoneal injection. Flow cytometric analysis of T cell proliferation and homing receptor expression on OT-I cells residing in lung at 45 days post MVA infection. Naïve OT-I Thy1.1^+^ cells were transferred into Thy1.2^+^ recipient mice one day before mice were infected with 1.8 × 10^6^ pfu MVA-Ova by s.s., i.t. or i.p.. At 45 days after infection, proliferation and tissue-homing receptor expression of OT-I T_RM_ cells isolated from lung tissue were analyzed by flow cytometry. Data are representative of three independent experiments (n = 5 mice per group). ESL, E-selectin ligand.

**Extended Data Figure 5.**
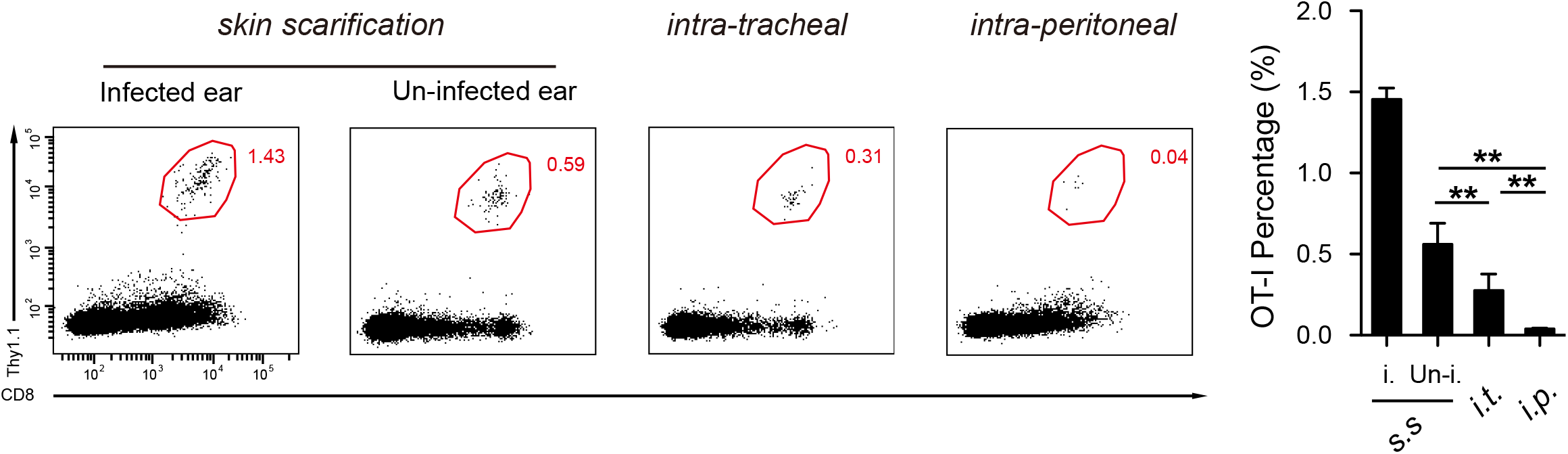
Skin T_RM_ cells generated by MVA infection via skin scarification, intra-tracheal administration or intra-peritoneal injection. Flow cytometric analysis and quantification of skin T_RM_ cells at day 45 post 1.8 × 10^6^ pfu MVA infection via indicated routes. Data are representative of three independent experiments (n = 5 mice per group). **p < 0.01.

## Online Methods

### Mice

Wide-type (WT) C57BL/6, CD45.1^+^, Thy1.1^+^, Rag1^-/-^, μMT, Langerin-DTA, Langerin-DTR mice were purchased from Jackson Laboratory. Thy1.1^+^ Rag1^-/-^ OT-I mice were maintained through routine breeding in the animal facility of Harvard Institute of Medicine, Harvard Medical School. Animal experiments were performed in accordance with the guidelines put forth by the Center for Animal Resources and Comparative Medicine at Harvard Medical School. Mice were randomly assigned to each group before start and experiments were performed blinded with respect to treatment. For survival experiments, mice that had lost over 25% of original BW were euthanized.

### Viruses

An attenuated strain (VACV) of WR-VACV was used in some experiments as control vaccine and was a kind gift from Dr. Bernald Moss (National Institutes of Health, Bethesda, MD). Wildtype WR-VACV were purchased from American Tissue Culture Company (ATCC). The virus stocks were expanded and tittered in Hela cells and CV-1 cells (ATCC) by standard procedures. ACAM3000MVA (Acambis Modified Vaccinia Ankara) (MVA) and DF-1 Cells were gifted by Dr. Michael Seaman (Beth Israel Deaconess Medical Center, Boston MA). MVA stocks were expanded and titrated in DF-1 cells as previously described (39, 40).

### Virus Infection

Mice were immunized with the MVA or VACV at the indicated doses by skin scarification as previously described. Alternatively, mice were immunized by s.c., i.d., or i.m. injection at the indicated dose. For secondary challenge, memory mice were challenged intranasally with a lethal dose of WR-VACV (2 × 10^6^ pfu in 20 μl of PBS) at 6 to 20 weeks post immunization. The change of BW and survival of mice were monitored daily following challenge for up to 12 days.

### *In vitro* restimulation assay

Poxvirus-specific T cell response against poxvirus was assessed at day 7 post challenge. Single cell suspension prepared from draining lymph nodes or spleens was re-suspended in T cell medium (RPMI containing 10% FBS, 2mM 2-β mercaptoethanol, 1X nonessential amino acid, 1X sodium pyruvate), and were used as effector cells. For target cell preparation, naïve splenocytes was infected at 37 °C for 5 h with WR-VACV at a MOI of 5 in RPMI medium supplemented with 10% FCS. After infection, the cells were washed 3 times with PBS, and cocultured (5 x 105 cells/well) with effector cell at a 1:1 ratio in 96 well plate in T cell medium at 37 °C for 48 h. Uninfected naïve splenocytes co-cultured with target cells were used as negative controls. IFN-γ concentration in the culture supernatants were measured by ELISA using anti-IFN-γ mAb pairs (BD Pharmingen) according to manufacturer’s protocol.

### Preparation of cell suspensions

Lymph nodes and spleens were harvested and pressed through a 70-μm nylon cell strainer to prepare cell suspensions. Red blood cells (RBC) were lysed using RBC lysis buffer (00-4333-57; eBioscience). Skin tissue was excised after hair removal, separated into dorsal and ventral halves, minced, and then incubated in Hanks balanced salt solution (HBSS) supplemented with 1 mg/ml collagenase A (11088785103; Roche) and 40 μg/ml DNase I (10104159001; Roche) at 37 °C for 30 min. After filtration through a 70-μm nylon cell strainer, cells were collected and washed three times with cold PBS before staining.

### Mouse adoptive transfer and treatment

Lymph nodes were collected from naïve female donor mice at age of 6-8 weeks. T cells were purified by magnetic cell sorting using a mouse CD8α^+^ T-cell isolation kit (130-104-075; Miltenyi Biotec) or a mouse CD4^+^ T-cell isolation kit (130-104-454; Miltenyi Biotec), according to the manufacturer’s protocols. T cells were then transferred intravenously into female recipient mice at a total number of 5 × 10^5^. T cells were labeled with carboxyfluorescein succinimidyl ester (CFSE, 65-0850; eBioscience) before co-transfer, where indicated. In some experiments, mice were treated daily with FTY720 (10006292; CAYMAN, 1 mg/kg) by intraperitoneal injection.

### Microarray, data analysis and quantitative real-time PCR

For each group of microarray dataset, OT-I cells from 15-20 mice were sorted with a FACSAria III (BD Biosciences) and pooled. RNA was extracted with a RNeasy Micro kit (74004; Qiagen). RNA quality and quantity were assessed with a Bioanalyzer 2100 (Agilent). Then RNA was amplified and converted into cDNA by a linear amplification method with WT-Ovation Pico System (3302-60; Nugen). Subsequently cDNA was labeled with the Encore Biotin module (4200-60; Nugen) and hybridized to GeneChip MouseGene 2.0 ST chips (Affymetrix) at the Translational Genomics Core of Partners Healthcare, Harvard Medical School. GeneChips were scanned using the Affymetrix GeneChip Scanner 3000 7G running Affymetrix Gene Command Console version 3.2. The data were analyzed by using Affymetrix Expression Console version 1.3.0.187 using Analysis algorithm RMA. To evaluate overall performance of microarray data, principal component analysis (PCA) and Pearson correlation coefficients among 12 diverse samples were applied by using 26,662 transcripts (R Program). All microarray data was submitted to the Gene Expression Omnibus.

For relative quantitative real-time PCR, RNA was prepared as described above. Bio-Rad iCycler iQ Real-Time PCR Detection System (Bio-Rad) was used with the following settings: 45 cycles of 15 s of denaturation at 95 °C, and 1 min of primer annealing and elongation at 60 °C. Realtime PCR was performed with 1 μl cDNA plus 12.5 μl of 2× iQ SYBR Green Supermix (BioRad) and 0.5 μl (10 μM) specific primers. For absolute quantitative real-time PCR. each standard curve was constructed using 10-fold serial dilutions of target gene template ranging from 10^7^ to 10^2^ copies per mL and obtained by plotting values of the logarithm of their initial template copy numbers versus the mean Ct values. The actual copy numbers of target genes were determined by relating the Ct value to a standard curve.

### Determination of viral load

Viral load in various tissues following MVA or VACV skin scarification was determined by quantitative real-time PCR, as previously described (17). Briefly, DNA was purified using the DNeasy Mini Kit (Qiagen, Valencia, CA). The primers and TagMan probe used in the quantitative PCR assay are specific for the ribonucleotide reductase Vvl4L of vaccinia virus. The sequences are: (forward) 5’-GAC ACT CTG GCA GCC GAA AT-3’; (reverse) 5’-CTG GCG GCT AGA ATG GCA TA-3’; (probe) 5’-AGC AGC CAC TTG TAC TAC ACA ACA TCC GGA-3’. The probe was 5’-labeled with FAM and 3’-labeled with TAMRA (Applied Biosystems, Foster City, CA). Real-time PCR was performed with the Bio-Rad iCycler iQTM Real-Time PCR Detection System (Bio-Rad Laboratories). Thermal cycling conditions were 50°C for 2 min and 95°C for 10 min for one cycle, followed by 45 cycles of amplification (94°C for 15 s and 60°C for 1 min). Standard curve was established from DNA of an MVA or VACV stock with previously determined titer. Corresponding CT values obtained by the real time PCR reactions were plotted on the standard curve to calculate viral load in the samples. The number of viral DNA copies was normalized to that in the skin samples of uninfected naïve mice.

### Statistical analysis

Comparisons for two groups were calculated using Student’s t test (two tailed). Comparisons for more than two groups were calculated with one-way analysis of variance (ANOVA) followed by Bonferroni’s multiple comparison tests. Two-way ANOVA with Holm–Bonferroni post hoc analysis was used to compare weight loss between groups and Log-rank (Mantel-Cox) test was used for survival curves. p < 0.05 was considered statistically significant.

